# *Kinex* infers causal kinases from phosphoproteomics data

**DOI:** 10.1101/2023.11.23.568445

**Authors:** Alexandra Valeanu, Verena Golz, David W. Avila, Manuel Tzouros, Juliane Siebourg-Polster, Laura Badi, Jitao David Zhang

**Affiliations:** Pharma Early Research and Development, Roche Innovation Centre Basel, F. Hoffmann-La Roche Ltd, Grenzacherstrasse 124, 4070 Basel, Switzerland

**Keywords:** cell signalling, phosphoproteomics, post-translational modifications, kinases, drug discovery

## Abstract

**Motivation:** *Phosphoproteomics data are essential for characterising signalling pathways, identifying drug targets, and evaluating efficacy and safety profiles of drug candidates. Emerging resources, including a substrate-specificity atlas and drug-induced phosphoproteomics profiles, may transform the inference of causal kinases. However, there is currently no open-source software that leverages insights derived from these resources.*

**Results:** *We introduce* Kinex, *a workflow implemented in the same-name Python package, which infers causal serine/threonine kinases from phosphoproteomics data. Kinex allows users to score kinase-substrate interactions, perform enrichment analysis, visualise candidates of causal regulators, and query similar profiles in a database of drug-induced kinase activities. Analysing seven published studies and one newly generated dataset, we demonstrate that analysis with Kinex recovers causal effects of perturbations and reveals novel biological insights. We foresee that Kinex will become an indispensable tool for basic and translational research including drug discovery.*

**Availability:** *Kinex is released with the GNU General Public License and available at https://github.com/bedapub/kinex.*

## 1 Introduction

The human genome encodes more than 500 kinases that recognise proteins as their substrates^1^. Kinase binds to its preferred substrates, partially by recognising the sequence motifs around a serine, threonine, or tyrosine residue, and attaches phosphate groups to the residue^2^. Protein phosphorylation plays a key role in signal transduction, which both mediates and is impacted by pathophysiological conditions^3,4^. Despite the importance, little is known about how kinase activities are modulated in diseases. New approaches to inferring kinase activities are warranted.

Kinases bear great potential as therapeutic targets for human diseases^5^. A total of 77 kinase-inhibitor drugs have been approved by FDA as of September 2023. However, development of kinase inhibitors as drug candidates is complicated by both limited understanding of functions of many kinases, and a lack of selectivity of many kinase inhibitors, which leads to unfavorable risk-benefit profiles due to excessive toxicity. Inhibitors approved so far for clinical use only target a limited number of kinases^6^ (Supplementary Table 1), and are applied in limited indications including oncology and autoimmune diseases. Beyond kinase inhibitors, other types of drugs exert their pharmacology and toxicology by regulating kinases directly or indirectly, too. New techniques to profile kinase modulation by chemical matters may catalyse the discovery of efficacious and safe kinase-modulating drugs.

Recent progress in the detection and quantification of phosphorylation through mass spectrometry has led to the identification of about 90,000 serine and threonine phosphorylation sites in the human proteome^7^. By employing a cell-free method known as the positional scanning peptide assay (PSPA), Johnson *et al*. created a quantitative atlas of intrinsic substrate specificity for 303 serine and threonine kinases^8^. Almost simultaneously, Zecha *et al*. reported a large-scale study of the post-translational modifications, comprising proteomics data of 31 drugs tested in 13 human cancer cell lines with varying time points and concentrations^9^. These data resources hold the promise to empower researchers better leverage phosphoproteomics data in at least three aspects. First, they may enable the inference of causal kinases from phosphoproteomics data derived from case-control experiments. Second, they may allow phosphoproteomics-based characterisation of primary and secondary targets of drug molecules. Finally, they may enable comparison of kinase-modulation profiles of new drugs with profiles induced by drugs with known mode of actions (MoA). To our best knowledge, however, there is currently no software tool that allows researchers to achieve the three goals.

To translate the potentials into practice, we introduce *Kinex* (*Kin*ome *ex*ploration), a workflow implemented in a Python package designed for the inference of causal kinases from phosphoproteomics data by integrating and leveraging multiple resources. Kinex takes differential phosphorylation data as input and infers activities of upstream, i.e. causal, kinases that most likely have led to the observed changes (Figure 1, panels a-g). We validated Kinex and demonstrated its capacity to resolve cryptic kinase-substrate relationships, to characterise MoA of drugs, to reveal the interplay of signalling pathways, and to analyse *in vitro* and *in vivo* mouse studies. Taken together, Kinex offers a new avenue to study cell signalling and to characterise therapeutic modalities such as small molecules, antibodies, macrocycles, etc.

**Figure 1:**
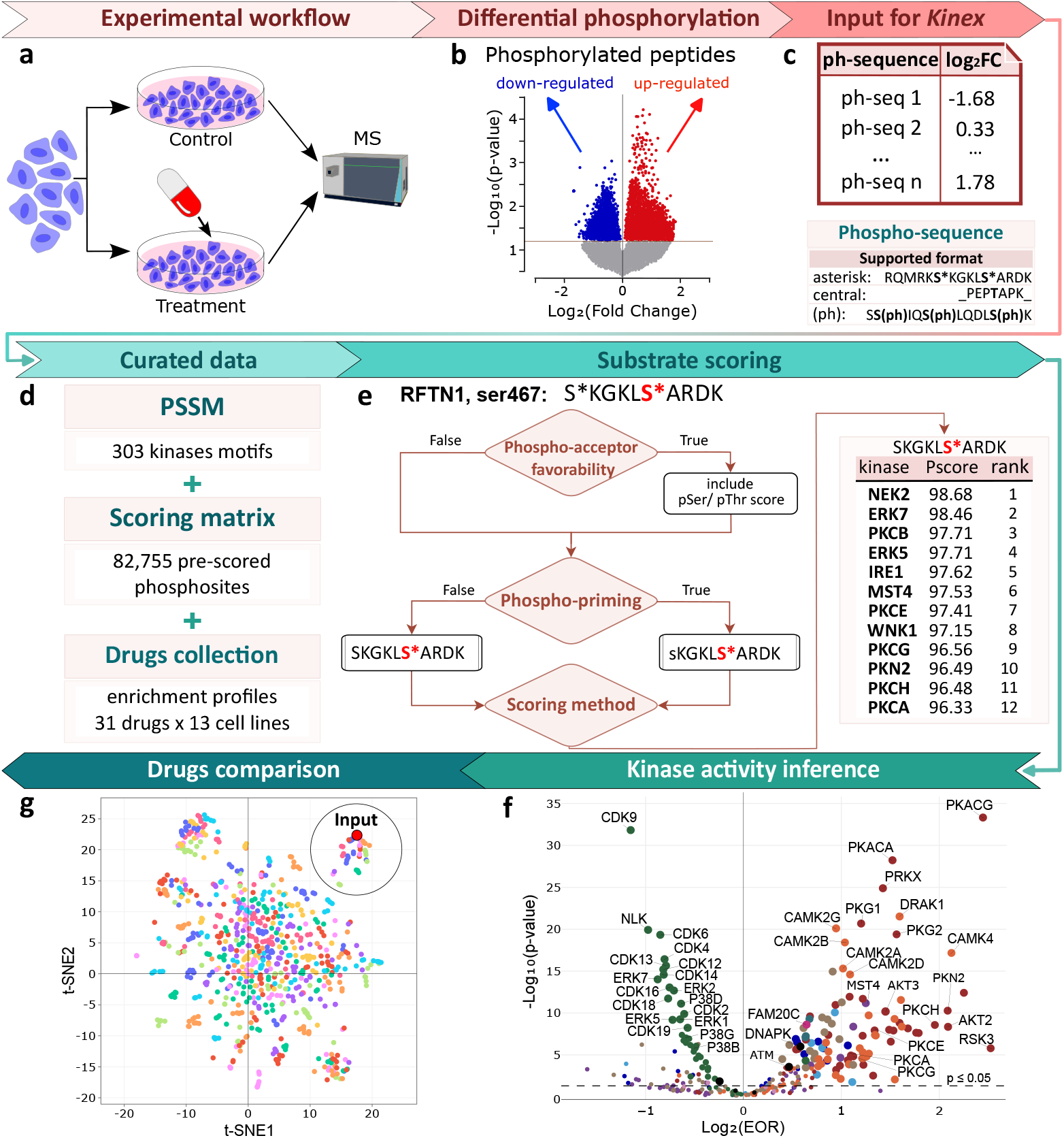
Phosphoproteomics data analysis workflow implemented in Kinex. **a** Kinex requires differential phosphorylation profiles as input, which can be derived from for instance case-control phosphoproteomics experiments. **b**,the volcano plot highlights differential phosphopeptides between treatment and control.**c**,Kinex requires a tab-limited file as input, which contains the modified phosphorylation sequences with one or multiple phosphorylation sites in the first column, and the log transformed fold changes in the second column.**d**, Kinex relies on two essential components for scoring substrates and inferring kinase activities: the Position Specific Scores Matrices (PSSMs) of 303 serine/threonine kinases, each scores 23 amino acids across 9 positions from -5 to +4, and a matrix of pre-calculated scores for the 303 kinases and 82,755 known human phosphosites. For the optional functionality of database query, Kinex requires the collection of drugs enrichment profiles which is freely downloadable.**e**, workflow and an exemplary output of the substrate scoring method. Kinex accounts for both phospho-acceptor favorability and for phospho-priming scores explicitly. The output shows the top scoring kinases that likely phosphorylates RFTN1 Ser467. In this example, phospho-priming was set to `False` and phospho-acceptor favorability to `True`.**f**, Inferred activity changes induced by 30,000 ng/mL of Rituximab in SU-DHL-4 cells after 2 hours treatment compared with vehicle control. Activated (*log*_2_(enrichment odds ratio, or EOR)>0) and inhibited (*log*_2_(EOR)<0) kinases are colored by kinase family membership derived from phylogenetic analysis. Selected kinases are highlighted with labels. The points above the dash line correspond to kinases which are significantly modulated (*p* ≤ 0.05). The analysis was run with a log2-transformed fold change threshold of 1 as threshold and the same aforementioned parameters.**g**, comparison of the Rituximab profile with all profiles derived from the *decryptM* platform. Each point represents a sample, which in this context means a unique combination of drug, concentration, the duration of the treatment, the cell line used, and the running index of replicate. Samples most similar to the input are highlighted by the enclosing circle. The origin point (0, 0) represents the effect of vehicle control, i.e. no changed kinase activities. The points are colored randomly to make the figure better readable.

## 2 Results

### Kinex is a comprehensive tool with new features

Quite a few software tools have been developed to identify kinases that may cause the observed changes in substrate phosphorylation levels^10^. We categorise them by methodology, the ability to score unknown substrates, and kinase coverage in Supplementary Table 2. Among them, *INKA*^11^, *RoKAI*^12^ and *Kinase Library*^8^ are of particular interest. Unlike other tools, they can handle unseen substrates, for instance synthetic peptides or sequences containing mutations. They are also able to infer most kinases: *RoKAI* and *Kinase Library* covers 261 and 303 protein kinases, respectively, while *INKA* can even infer unknown kinases.

Kinex retains the flexibility with unseen substrates and the wide coverage of kinases, while offering four additional key functionalities. First, the statistical model of Kinex considers phospho-priming explicitly. This is critical because sequences with multiple phosphorylation sites account for about 12% of all peptides in a typical phosphoproteomics dataset, and we observed that modelling them explicitly increases the sensitivity and power of inference (Supplementary Figure 1). Second, Kinex does not only support central and asterisk formats of sequences as input, but also the modified sequence format produced by *MaxQuant*^13^, a widely used open-source software for mass-spectrometry data analysis. Third, Kinex offers a database of inferred causal kinase profiles induced by drug treatments derived from the *decryptM* platform. The query functionality allows users to interpret kinase activity changes in the context of pharmacological perturbations. Finally, implemented as a open-source Python package, Kinex allows all users to analyse large-scale phosphoproteomics data with command-line and integrate the functionality into their own computational workflows, instead of having to submit data to a web page. Taken together, these novel features make Kinex a user-friendly and powerful tool that complements existing algorithms.

### Example applications

We first validated the core ability of Kinex to recover individual perturbed kinases with published studies (Supplementary Methods). Next, we applied Kinex to a number of case studies to demonstrate its value for cell-signaling research and drug discovery.

#### Unveiling kinase-substrate relationships

Kinex can shed light on cryptic kinase-substrate relationships. We take the example of RFTN1, the lipid raft linking protein, to illustrate the point. RFTN1 can be phosphorylated at mulitple sites. Among them, phosphorylation of the site Ser467 plays an important role in both B-cell antigen receptor (BCR) signalling and the maintenance of lipid raft integrity^14^. The kinases responsible for phosphorylating this site remain elusive^9^.

We reasoned that analysing the dose- and time-dependent phosphoproteomics datasets derived from the *decryptM* platform^9^ with Kinex may offer new insights. To this end, we performed differential phosphorylation analysis of RFTN1’s Ser467 site with data generated from human SU-DHL-4 cells (a cellular model of human B-cell lymphoma) treated with Rituximab, a monoclonal antibody targeting the CD20 receptor^9^, and applied Kinex to infer causal kinases regulating the site.

Compared with time-matching controls, Ser467 phosphorylation is mostly positively regulated up to 2 hours post treatment, though the induction first increases and then decreases along time. Critically, the effect size is correlated with the concentration of Rituximab (Supplementary Figure S3, panel a and b), and the regulation is specific to this phosphorylation site (Supplementary Figure S3, panel c). Taken together, the site-specific, time- and concentration-dependent profile suggests that Rituximab is likely to modulate RFTN1 Ser467 phosphorylation indirectly via kinases downstream of CD20.

We ranked kinases inferred by Kinex by their propensity to phosphorylate the Ser467 site: NEK2, a member of the NEK kinase family, ERK5 and ERK7, two kinases of the ERK family, and several members of the protein kinase C (PKC) family are most likely upstream kinases (Figure 1e). Manual inspection confirms that the sequences flanking the site match the motifs recognised by these kinases the best. Intriguingly, the inferred activities of these kinases show contrasting time dynamics: while inferred activities of NEK2 and PKCs show a highly correlated temporal profile with that of RFTN1 Ser467 phosphorylation, the inferred activities of ERK5/ERK7 are partially anti-correlated (see Figure 1f for inferred results after 2h, and Supplementary Figure S3, panel d for the complete time course). The results suggest that RFTN1 Ser467 phosphorylation may be regulated by multiple kinases with distinct temporal activity profiles.

The results raise the intriguing hypothesis that RFTN1 Ser467 phosphorylation may be the convergent point of multiple signalling and metabolism pathways. NEK2 is often found up-regulated in diffuse large B cell lymphoma, an indication of Rituximab. NEK2 cooperates with the Hippo pathway^15^, affects glycolysis^16^, and affects mouse B cell development by involving in multiple pathways including the AKT pathway and TGF-*β* pathway^17^. The ERK/MAPK pathway constitutes the signalling cascades downstream of BCR signalling together with PI(3) kinase, NF-AT, and NF-*κ*B pathways^18^. Protein kinase C (PKC) family members play important and distinct roles in B-cell activation, in particular B-cell survival and NF-*κ*B pathway activation following BCR signalling activation^19^. While these pathways have been individually described to affect B-cell biology, the analysis with Kinex allows us to speculate that they may all regulate RFTN1 to affect lipid raft integrity and BCR signalling. Further research is warranted to explore the value of RFTN1, and more specifically the Ser467 site, as a novel biomarker or therapeutic target for B-cell lymphoma.

In summary, Kinex suggests that NEK2, ERK5, ERK7, and several PKC-family kinases are likely candidates for causal kinases phosphorylating RFTN1. The case study shows the capacity of Kinex to shed light on cryptic kinase-substrate relationships.

#### Revealing mode of action of drugs

We speculated that Kinex can be used to characterise the MoA of drugs, in particular their impact on human kinase activities. To explore the potential, we expanded the case study above, and examined kinome-wide results derived from human lymphoma cells SU-DHL-4 treated with Rituximab^9^.

Kinex inferred that Rituximab affects many kinases in a time- and concentration-dependent manner. Focusing on 2 hours post treatment with the highest concentration, we observed that Rituximab regulates multiple kinases associated with cell cycle and apoptosis. In particular, it inhibits cyclin-dependent protein kinases, which mediate the growth inhibitory cues of upstream signalling pathways^20^. Also, it promotes apoptosis by inhibiting multiple kinases belonging to the ERK and MAPK pathways^21,22^(Figure 1f). The results corroborate the phenotypic data reported by Zecha *et al*.

We wondered whether treatment with other drugs can induce a similar profile with that of Rituximab. With the query functionality of Kinex, we looked up the most similar profiles of upstream kinases, and found that the five top hits were all associated with Rituximab, though the cell line, exposure time, and dose vary (Figure 1g). It suggests that Rituximab has a unique MoA among the drugs tested by Zecha *et al*.^9^. Indeed, Rituximab was the only antibody-based drug targeting CD20: the only other two antibody-based drugs both target HER2 (Pertuzumab and Trastuzumab), and none of the other 30 drugs has CD20 as the primary or secondary target. If another CD20-targeting drug was tested, we would have expected some similarity with the profile of Rituximab. This was, in fact, the case for Pertuzumab and Trastuzumab, two antibodies targeting HER2 with distinct mechanisms of actions. Inferred kinase activity changes induced by both antibodies are partially correlated between 10ng/mL Pertuzumab and Trastuzumab after 2h treatment in BT474 cells (Pearson’s correlation coefficient *ρ* = 0.44).

In short, researchers can use Kinex to characterise MoA of drugs, and to compare drugs with regard to their modulation of kinase activities.

#### Dissecting interplay of signalling pathways

Having applied Kinex successfully to single perturbations, we wondered whether we can use it to dissect the interplay between two signalling pathways. we chose the Angiopoietin-2 (Ang-2, or Ang2) pathway and the Vascular Endothelial Growth Factor A (VEGF-A, or VEGF) pathway as a case study.

Both Ang2 and VEGF are growth factors critically involved in multiple biological processes, in particular the homeostasis of endothelial barrier permeability and angiogenesis^23,24^. The effect of Ang2, like other angiopoietins (Ang1, Ang3, and Ang4), is mediated through the tyrosine kinase receptors Tie1 and Tie2^25^. The effect of VEGF is mediated via VEGF receptors (VEGFRs), including VEGFR1 (also known as FLT1), VEGFR2 (KDR), and VEGFR3 (FLT4)^26^. Though the effect of Ang2 and VEGF on endothelial barrier permeability has been well studied, the interplay of the two pathways remains little understood.

To address the challenge, we generated a new phosphoproteomics dataset using human umbilical vein endothelial cells (HUVECs) as an *in vitro* model of endothelial barrier permeability. HUVECs were treated with Ang2 (300 ng/mL), VEGF (5 ng/mL), and the combination of Ang2 and VEGF (100 ng/mL and 5 ng/mL, respectively) for 20 minutes, followed by a phosphoproteomics workflow (Figure 2a and Supplementary Methods). We performed differential phosphorylation analysis and fed the output to Kinex to infer kinases modulated by Ang2, VEGF, and by their combination.

**Figure 2:**
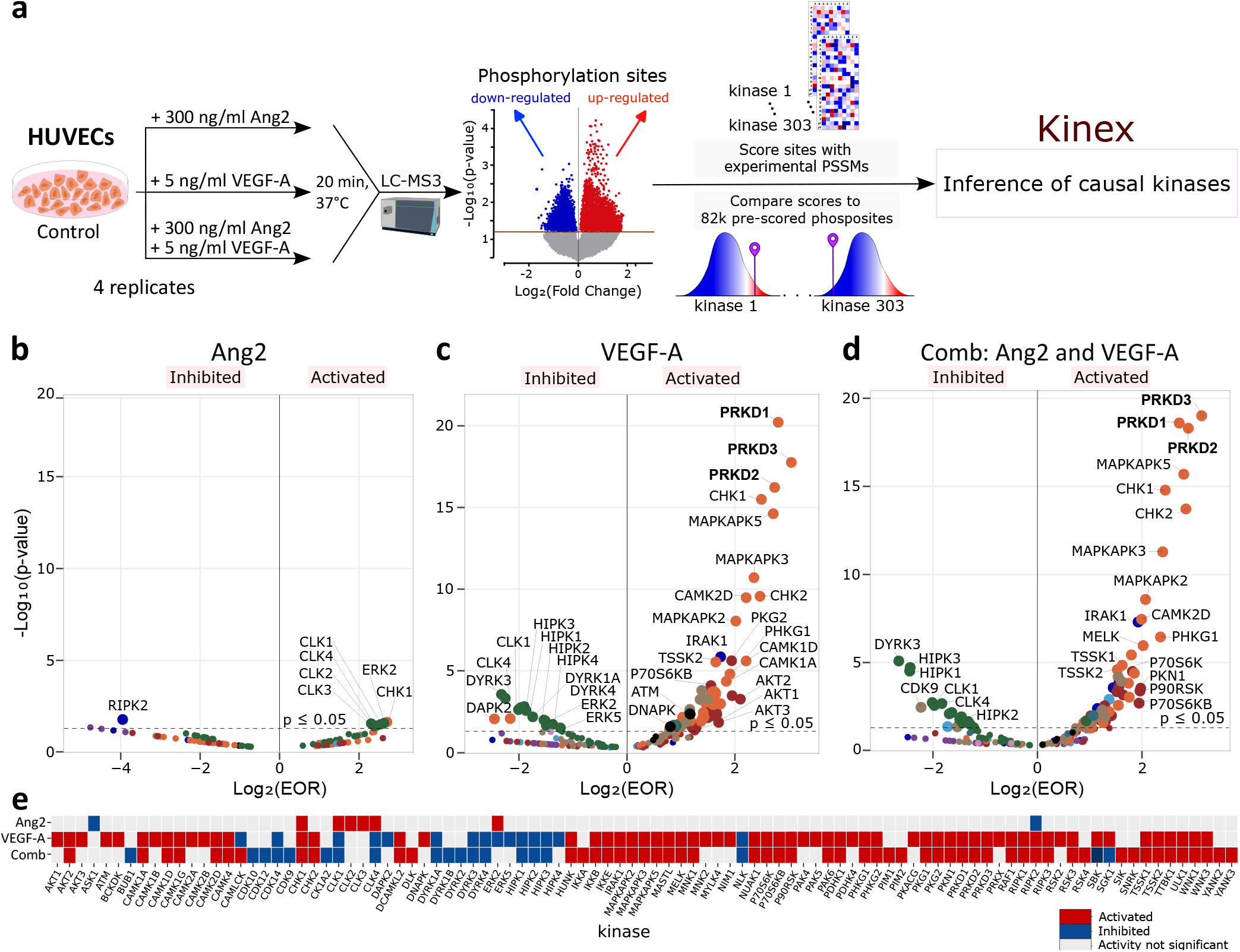
Effect of Angiopoietin-2 signalling, VEGF-A signalling, and their combination on kinase activities in HUVECs. **a**,Schematic of the experimental and data pre-processing workflow. Differential phosphorylation data was used as input for Kinex.**b**,**c**,**d** inferred changes of kinase activities upon treatment of Ang2, VEGF-A, and the combination. The inference analysis was conducted with phospho-acceptor favorability set to `True` and a log2-transformed fold change threshold of 1. The points above the dash lines correspond to kinases with statistically significant changes (*p* ≤ 0.05). The kinases are colored by their phylogenetic relationship.**e**, a heatmap illustrates the significantly modulated kinases (in columns) across all three treatment conditions (in rows). Red, blue, and gray cells indicate significant activation, significant repression, and non-significant changes (*p*>0.05), respectively.

Analysis with Kinex revealed that both Ang2 and VEGF signalling modulates multiple kinases. While the effects of Ang2 are restricted to a few kinases including cyclin-like kinases CLKs, ERK2, CHK1, and RIPK2, VEGF affects a large number of kinases (Figure 2, panels b-d). Both single treatment with VEGF and its combination with Ang2 were inferred to activate virtually all kinases of the protein kinase D (PKD) family. It has been well established that the binding of VEGF to VEGFR2/KDR leads to rapid and strong activation of PRKD (Protein Kinase D) in HUVECs^27,28^, which has profound consequences on gene expression^29,30^. At the same time, we observed that VEGF represses the activity of several HIPKs repressed. HIPK1 and HIPK2 were reported to act as negative regulators of VEGF. Their loss leads to up-regulation of *MMP10* and *VEGF* genes, causing excessive proliferation of endothelial cells and poor junction formation^31^.

Notably, the effect of the combo treatment resembles more the effect of VEGF than a linear sum of the effect of individual treatments (Supplementary Figure S4). This observation suggests that in the experimental condition, kinase-level regulation by the combination of VEGF and Ang2 is primarily dominated by the VEGF signalling pathway. At the same time, we observe a number of kinases for which the inferred activity differs significantly from the profile upon VEGF-A treatment, including CLK3, BCKDK, SBK, and AKT3 (showing repressed activity in the combo treatment but increased activity in VEGF treatment), as well as DAPK2, ERK2, ERK5, CAMLCK, PIM2 (showing higher activity in the combo treatment compared with in the VEGF treatment). In the context of renal cell carcinoma, increased activity of Branched-chain keto-acid dehydrogenase kinase (BCKDK) promotes vascular permeability and angiogenesis of HUVECs^32^. AKT3 regulates VEGF, and intravitreal delivery of AKT3 siRNA in the diabetic rat significantly decreased the expression of both AKT3 and VEGF^33^. ERKs are found to be among kinases involved in temporally-resolved kinase regulatory networks that control endothelial barrier integrity^34^. Furthermore, a few other kinases such as ATM, CAMK2A/B, and PAK4/5 show different profiles between VEGF and the combo (Figure 2e). While the consequence of regulation of these and other kinases remain to be verified, simultaneous activation of VEGF and Ang2 pathways apparently does not simply pheno-copy the effect of VEGF. Instead, kinases involved in permeability such as BCKDK, AKT3, and ERKs are modulated by the presence of Ang2.

Taken together, phosphoproteomics profiling of HUVECs treated with VEGF, Ang2 and their combination suggests that VEGF signalling has a broad impact on the Ser/Thr kinase network, while Ang2 signalling specifically modulates a few kinases, especially cyclin-like kinases. When both pathways are activated simultaneously, the outcome is largely dominated by the effect of VEGF. Intriguingly, a few kinases associated with endothelial cell permeability show different kinase activity profiles compared with VEGF treatment alone. It suggests that activation of Ang2 pathway may modify behavior of HUVECs when VEGF is present. Further research is warranted to explore the effect of drugs targeting both pathways in a similar setting, and to establish the relevance of these findings in clinical studies.

#### Applicability to mouse studies

Since all examples above use human data, we further challenged Kinex with phosphoproteomics data from mouse cell lines and animal studies, given their great importance both in basic and applied research as well as lines of evidence supporting such a challenge (Supplementary Methods).

First, we curated and analysed a phosphoproteomics dataset of mouse C2C12 myoblasts. The researchers generating the dataset deleted both catalytic sub-units, *α* and *α*’, of the protein CK2 (casein kinase 2) genetically, compared them with wild-type cells with phosphoproteomics, and characterised the knock-outs with a number of techniques^35^.

The team reported data from two clones (clone A and clone B) used in the genetic knockout experiment. Thus we were able to perform differential phosphorylation analysis for both clones, compare knock-outs with wild-type cells, and use the differential phosphorylation profiles as inputs for Kinex to assess the causal kinase in two clones independently. As expected, Kinex successfully identified both catalytic sub-units of CK2 as the kinases whose activities are most significantly inhibited (Figure 3a). This corroborates authors’ claim that they have generated two specific CK2 knockouts.

**Figure 3:**
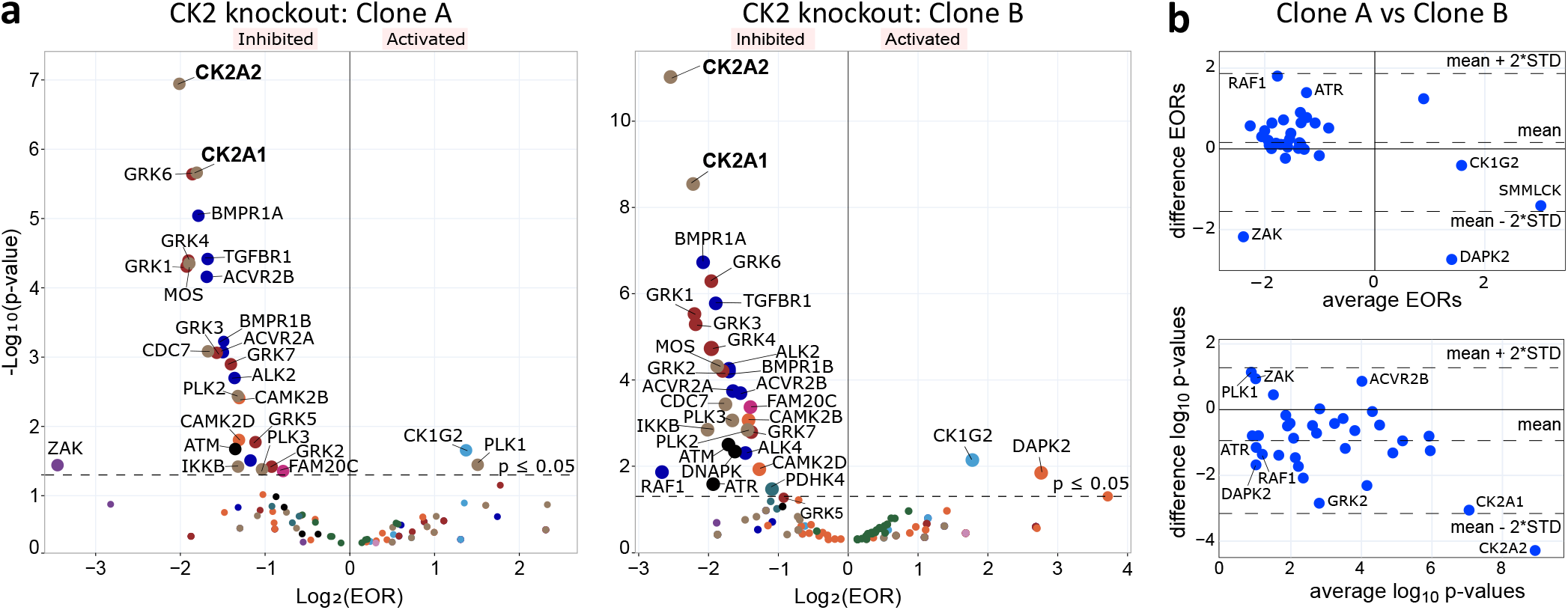
Recovery of CK2 knockout in mouse C2C12 cells by Kinex. **a**, inferred kinase activity changes in two cell clones, A and B, 10 min after genetic deletion of the CK2 catalytic sub-units.**b**,Comparison between clone A and B using the significantly modulated kinases (*p* ≤ 0.05) and the MA plot^1^. In the MA plot, each dot represents one kinase. The arithmetic mean of inferred activity changes in two clones compared with vehicle control is plotted on the x-axis, while difference between the two are plotted on the y-axis. Solid lines indicate *x* = 0 and *y* = 0, respectively. Dash lines indicate twice the standard deviation of all kinases and assist spotting outliers.

We assessed the agreement of inferred kinase activities between the two clones (Figure 3b). Interestingly, besides CK2, we observed other kinases with consistent changes of inferred activities in both clones. They include kinases involved in cell communication (Grk1-7)^36,37^, cell growth and proliferation (ALK2,4 and ACVR2*α*/*α*’)^38^, osteogenesis (BMPR1*α*/*β*)^39,40^, DNA damage and stress response (ATM, ATR and DNA-PK)^41,42^, and cell-cycle control (PLK1,2,3)^43,44,45^. These results suggest that genetic knockout of CK2 catalytic sub-units disrupt multiple cellular pathways, consistent with previous reports on the pleiotropic roles of CK2^46^,

Intriguingly, besides overwhelmingly consistent downstream changes, we also observed considerable difference of other kinases, for instance ZAK (sterile alpha motif and leucine zipper containing kinase AZK) and DAPK2 (death-associated protein kinase 2). Apparently, the two biological replicate caused slightly different activities of some kinases. We are uncertain whether off-target effects of the CRISPR/Cas9 protocol would explain the difference. Nevertheless, our observations suggest that beyond being able to detect primary and secondary causal kinases, Kinex can detect subtle variability between biological or technical replicates, and thus is useful for quality-control purposes.

In summary, the CK2 example highlights the capability of Kinex to infer primary causal kinases with experiments in mouse cells, to reveal consequent events caused by the primary perturbation, and to assist quality control. Can we use Kinex also with *in vivo* phosphoproteomics studies? If so, this may open up new avenues to study signalling pathways and MoA of drugs in physiologically relevant conditions.

To answer the question, we next probed with Kinex an *in vivo* phosphoproteomics dataset, which consists of a time-series experiment conducted in mouse liver after insulin treatment^47^. The study reports “early” (5, 10, 15, 30 seconds) and “intermediate” (0.5, 1, 3, 6, 10 minutes) stages of insulin signalling in mouse liver after *in situ* hepatic insulin stimulation of fasted mice.

Applying Kinex to differential phosphorylation profiles with time-matching samples, we identified a large number of kinases with inferred activity changes. In order to illustrate the most important events in the kinase network, we selected kinases with the strongest enrichment, either with regard to effect size or with the statistical significance, from each time point, and capture their dynamics by visualising their enrichment values in a heatmap (Figure 4a).

**Figure 4:**
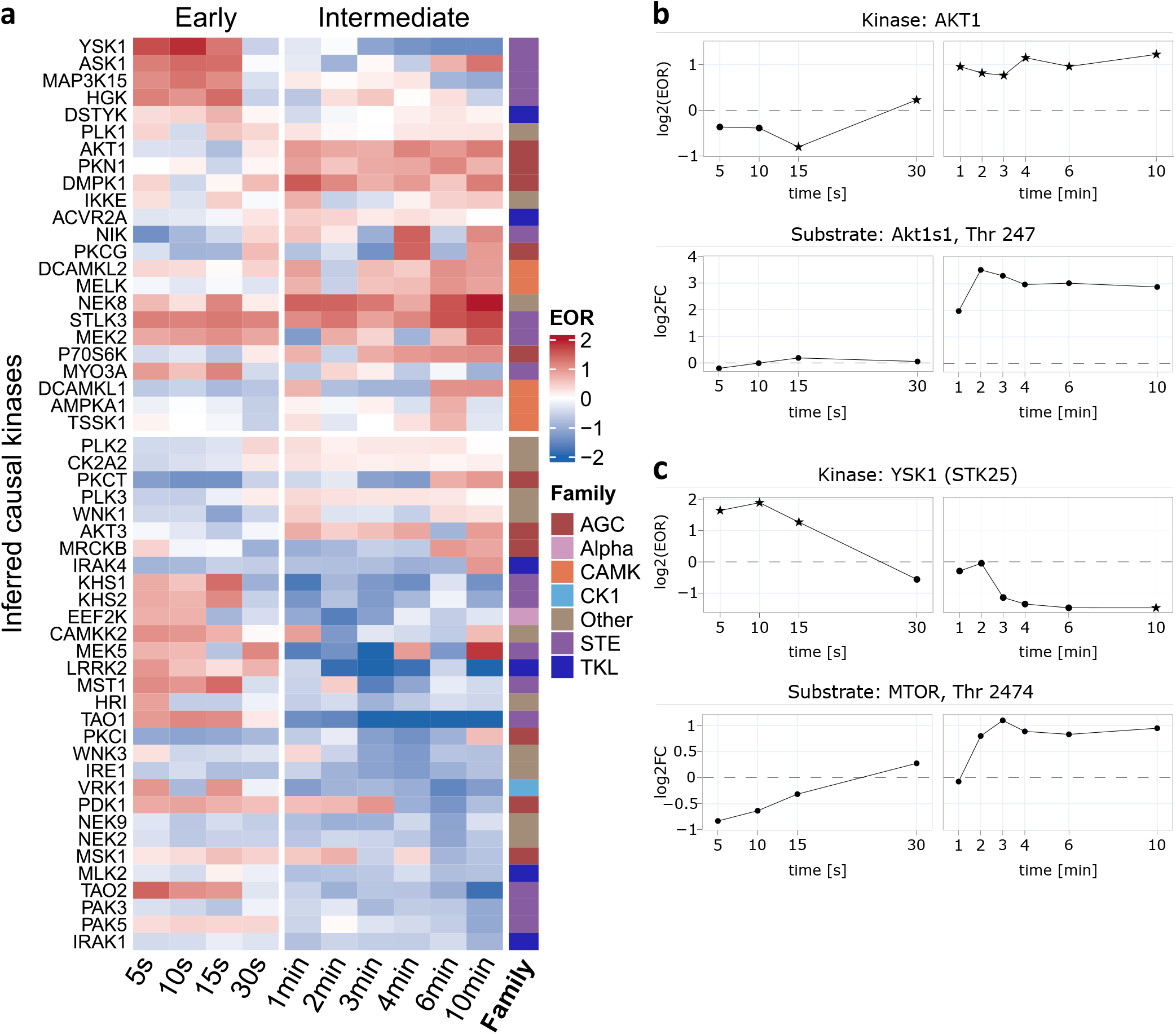
Kinase activity in early and intermediate insulin signalling. **a**, a heatmap of most prominently modulated kinases. Each row represents one kinase, which is colored to the right by the family membership. Each column indicate one time points, which are categorised into early and late phases^2^. We selected four kinases from each time point: the activated and repressed kinases with the smallest *p* values, or with the largest absolute enrichment odds ratio (EOR) values. Kinases that were selected in multiple time points are shown only once. The color gradient and intensity in the heatmap represents the enrichment odds ratio (EOR) score, with dark blue and red indicating repressed and increased activity, respectively. The top/bottom split represent the maximum amplitude of regulation across the time-series: strongest regulation of kinases in the top half is positive, while that of kinases in the bottom half is negative. The log2-fold threshold for the analysis was set to 1.5 in order to be consistent with the original publication. Since both the authors and we observed discrepancy in data of “early” 30 seconds and “intermediate” 0.5 minutes, we averaged the inference results of the two time points .**b**, time-dependent changes of inferred activities of the AKT1 kinase, and concurrent changes in the phosphorylation levels of its established target Akt1s1 at the phosphosite T247.**c**,, time-dependent changes of inferred activities of YSK, and concurrent changes in phosphorylation levels of MTOR at phosphosite T2474. Dash lines indicate *x* = 0; asterisks: *p* ≤ 0.05; circles: *p >* 0.05.

We observed intricate patterns of kinase activity changes with time. Within 5-15s, STE kinases seem to get activated, for instance YSK1, ASK1, HGK, and KHS1/KHS2. For most of them, the activity decreases after 1 min, suggesting that their activation is transient and is likely under tight control. Later, between 1 and 10min, members of the AGC family, for instance AKT1 (Figure 4b) and DMPK1, as well as members of the STE and CAMK families, such as MELK and MEK2, respectively, are activated. At the same time, a few kinases such as PAK3 and IRAK1 are suppressed. The majority of them are known to physically interact with each other, or considered to be relevant in literature, as revealed by an analysis with the STRING database^48^ (Supplementary Figure S6). A large number of proteins reported by Kinex belong to AKT/PKB (protein kinase B), MAPK, and AMPK (AMP-activated protein kinases) subfamilies, which are known to be integral components of the insulin signalling pathway^49^.

We also noticed significant modulations of other kinases that have not been extensively studied in this context. For example, TAO (thousand-and-one amino acids) kinases TAO1 and TAO2 are both activated early and then repressed at later time points. Their profiles are very similar to other kinases in the mammalian STE20-like (MST) protein kinase family, including YSK1 (Figure 4c), MYO3A, MST1, KHS1, KSH2, PAK3, and PAK5^50^, all of which were picked up by Kinex. STE-family kinases regulate the MAPK pathway and plays an essential role in insulin signalling and energy metabolism. Qi *et al*.^51^ profiled in-vivo kinase activity in human skeletal muscle from lean controls and adults with obesity and insulin resistance. STE was the largest family among the 54 active kinases they identified. In the context of insulin signalling *in vivo*, however, the roles of TAO1 and TAO2 remain elusive and warrant further study.

In summary, Kinex can be applied to phosphoproteomics data derived from *in vitro* and *in vivo* mouse experiments to infer causal kinases.

## 3 Discussions

Kinex infers causal kinases from phosphoproteomics data. We demonstrated the capacity of Kinex to uncover kinase-substrate relationships, to characterise MoA of drugs, to reveal the interplay of signalling pathways, and to analyse *in vitro* and *in vivo* data generated from mouse models. Available as an open-source software, Kinex empowers academic and industrial, basic and applied research alike.

Beyond the apparent value of Kinex for signalling transduction and kinase biology, we foresee that Kinex has the potential to update the standard practice of compound profiling and MoA studies in drug discovery. State-of-the-art methods include *in vitro* assays with the catalytic domain of kinases, for instance the CEREP/DiscoveryX panels^52^, the Kinobead assay, which profiles the affinity of kinases in cell lysates^53,54^, as well as the Nanobret assay, a cellular assay probing more than 200 recombinantly overexpressed kinases^55^. Emerging ABPP (activity-based proteomics profiling) techniques hold great promises, too (Zhao *et al*.^56^ and Gagestein *et al*., manuscript in preparation). While all these methods focus on the binding event of drug molecules to protein (kinase) targets, Kinex reveals uniquely the functional consequences of direct binding and indirect regulation of kinases *in vitro* or *in vivo*. Kinex expands the toolbox for drug profiling.

Together with phosphoproteomics studies, Kinex has the potential to be applied across the value chain of drug discovery and development. As the CK2 example suggests, it can be used to assess the downstream effects of modulating new drug targets. In the same vein, as the Rituximab example highlights, Kinex may be used to characterise the MoA and safety profile of lead molecules and drug candidates, thereby prioritising drug candidates for further development. The Ang2/VEGF example shows that Kinex may support developing combo therapies. And the ability of Kinex to analyse *in vivo* data makes it an attractive tool to derived valuable insights from animal studies for pharmacokinetics, pharmacodynamics, and safety evaluation. Furthermore, we foresee that it may become a cornerstone of multi-modal characterisation of human biology models including emerging platforms such as organoids and organ-on-a-chip, and be used to evaluate their alignment with human biology. The Kinex workflow can be used jointly with other technologies, such as transcriptional profiling, including single-cell RNA-sequencing and targeted approaches such as molecular phenotyping^57,58^, epigenetic profiling, proteomics, and metabolomics to construct the causal chain of drug’s effect. Furthermore, Kinex can be used to re-purpose existing drugs and to explore tumor- or individual-specific optimal treatment regiments, as suggested by a proof-of-concept study conducted with the INKA tool^59^.

While Kinex has enhanced our capability to analyse phosphoproteomics data, it has a few limitations in its current form. First, it is not yet capable to infer upstream tyrosines due to the lack of a tyrosine kinase substrate atlas. Though the situation may change soon^8^, inferring tyrosine-phosphorylating kinases can be challenging given the short half-life of phosphorylation events and complex regulations by constitutive and regulated phosphatases^60^. A similar limitation applies to non-protein kinases, too. Second, we could not run the Kinex for some peer-reviewed phosphoproteomics studies, because it requires a background list of peptide sequences (i.e. un-regulated sites), which are not always provided by publications. We recommend that research groups share the complete dataset to help others reproduce their findings. Third, currently Kinex does not model the regulation of protein half-life by phosphorylation events explicitly^61,62^, which may bias differential phosphorylation analysis. Finally, we have been unable to identify studies where the same perturbation is applied to both human and non-human cells. Such data are critical to demonstrate the specificity and sensitivity of across-species inference.

As phosphoproteomics techniques are progressing fast, and given the essential role of kinase in human biology and disease, we hope that Kinex’s functionality, accessibility, and capacity for comparative analysis make it a useful tool for researchers.

## 4 Methods

To build Kinex, we downloaded, curated, and integrated experimental data from multiple resources^1,8,9^ and implemented a novel algorithm to infer causal kinases. It takes sequences of phosphorylated peptides and changes in phosphorylation level as input, and reports a ranked list of kinases by the inferred activity of upstream kinases. Besides the linear motif, it models the presence of multiple phosphosites explicitly. Optionally, users can query similar profiles of drug-induced kinase activity changes. Details are provided in the Supplementary Methods.

## Data availability

The table of pre-scored peptides containing serine and/or threonine phosphorylation sites for 303 Ser/Thr kinases are available at https://zenodo.org/doi/10.5281/zenodo.10201141. All other data are included in the Kinex package, or in the supplementary information and tables.

## Acknowledgements

We thank Johnson *et al*., Zecha *et al*., and other teams who made their data accessible. We are especially indebted to Florian Bayer for his extensive help with pre-processing the *decryptM* data. We received critical feedback from Uwe Grether, Arne Rufer, Melanie Guerard, Andreas Zeller, Wolfgang Haap, Christian Kramer, Martin Ebeling, Bjoern Titz, Cheikh Diack, members of the Bioinfo Club, the Small-Molecule Data Analytics Network, and the Predictive Modeling and Data Analytics (PMDA) chapter led by Fabian Birzele. A.V. was supported by the Roche Advanced Analytics Network (RAAN). V.G. was supported by Roche Postdoc Fellowship (RPF).

## Author and Contributions

JDZ designed the study and supervised the work. AV wrote the code and performed the analyses. VG, MT and DWA designed and performed the phosphoproteomics experiment with Ang2 and VEGF in HUVECs; JDZ, JSP, LB and AV analysed the data. AV and JDZ generated the figures. AV and JDZ wrote and edited the manuscript with input from all of the authors. All authors read and agreed to the final manuscript.

