## Supplementary Methods for "*Kinex* infers causal kinases from phosphoproteomics data"

#### Analysis of kinase inhibitors approved by FDA

We curated the list of kinase inhibitors that are approved by the U.S. Food and Drug Agency (FDA) for clinical use as of 19. September 2023, assessed from www.brimr.org on November 11, 2023. We annotated their pharmacological targets and measurements of affinity or functional inhibition. In addition, we annotated pharmacodynamic studies performed with these drugs in mouse or rat models (excluding xenograft models) with regard to the primary targets, using assessment reports released by FDA and European Medicines Agency (EMA) of the drugs as well as publications. We classify the relationship between the kinase inhibitor and the mouse orthologue of the human kinase target of the drug is likely conserved when one of the two criteria is met: (1) the affinity or functional inhibition of a drug is tested both against its human kinase target and the mouse orthologue of the target, or (2) the pharmacokinetic-pharmacodynamic (PK/PD) relationship has been established in at least one study in mouse and rat.

We make two notes about our definition of conserved relationship. We excluded xenograft models since they may represent human kinase-inhibitor relationships, not those specific to mouse. Pharmacokinetic and pharmacodynamic (PK/PD) studies in rat were considered as an indicator for conserved kinase-inhibitor relationship, because bioinformatics analysis revealed that human, mouse, and rat genome contain more than 500 common and well behaved orthologues, and no rat kinase is shared with human only but not with that of mouse, which suggests no loss of rodent kinases in the mouse lineage.

Among 77 kinase inhibitors that are approved by FDA and/or EMA, we found 49 (64%) inhibitors likely target the mouse orthologue(s) of the human kinase target. Since the majority of the rest 28 inhibitors target mutated kinases, they are not always tested against the corresponding mouse mutants, or at least the data is not available.

#### Data provided by Kinex

(i) **Position Specific Scores Matrix (PSSM):** Kinex offers normalised and scaled scores from PSPA data of 303 human serine/threonine kinases and 23 amino acids (20 natural amino acids, plus phospho-serine, -threonine, and -tyrosine) across 9 positions (from -5 to +4, excluding the phosphorylation site) as seen in Supplementary Table 5 and Supplementary Figure 7. Additional columns for serine and threonine phospho-acceptor favorability are incorporated.

(ii) **Scoring matrix:** Kinex offers a reference table containing 82,755 pre-scored peptides containing serine and/or threonine phosphorylation sites for 303 kinases.

Kinex computes position-amino acid specific scores extracted from the PSSM table and returns the aggregated score for each kinase. Kinex then assesses the likelihood of the sequence being targeted by each kinase using the logarithm-transformed score in relation to the reference-scoring matrix.

#### Substrate scoring by Kinex

Kinex requires the sequence of amino acids. No additional information such as protein accession keys or gene names are needed. It can score any substrate, for which upstream kinases are known or not, as long as it has a serine or threonine as phospho-acceptor. For a given substrate, the score for each kinase is the product of the amino-acid scores in the corresponding positions. The amino-acid position scores are found in the normalized and scaled position specific scoring matrix (PSSM).

#### The phospho-priming effect

Johnson *et al.*[1] used a combinatorial peptide assay to score each amino acid including phosphoresidues (threonine/tyrosine). For serine phosphoresidue they took the same score as for the threonine one. In the PSSM matrix, the phosphoresidues are denoted with lowercase letters (s/t/y).

A phosphorylated residue within a peptide can have complex effect on the modification of other sites. The effect, known as phospho-priming, is prominent for kinases which recognise previously phosphorylated residues. Phospho-priming leads to non-linear and often non-trivial effect on the phosphorylation of their adjacent sites[2]. Some kinases groups such as CK1 (casein kinases 1) even exhibit a higher preference for sites with nearby phosphorylated residues (Supplementary Figure 8).

For example, if we have a multisite sequence **S(ph)**KGKL**S(ph)**ARDK and if we wish to include the phospho-priming, the software splits the sequence as follows:

a. **S(ph)**KGKL**s**ARDK: This keeps the first phosphorylation site and computes the score around it, while making the other sites lowercase.
b. **s**KGKL**S(ph)**ARDK: This keeps the second phosphorylation site and computes the score around it, while making the other sites lowercase.

The final score for **S(ph)**KGKL**S(ph)**ARDK is either:

1. `min(a,b)`, which returns the minimum percentile score between the sites for each kinase
2. `max(a,b)`, which returns the maximum percentile score between the sites for each kinase
3. `avg(a,b)`, which returns the average percentile score between the sites for each kinase
4. `all`, which returns the split sequences. Note that this option is not available for inferring kinase activity.

There is no limit to the number of phosphorylation sites within a phosphopeptide. By encoding amino acids all in uppercase, the explicit modelling of phospho-priming is disabled.

Peptides containing multiple phosphorylation sites account for on average 12% of all peptides in a typical dataset (Supplementary Figure 2, panel a), and with experimentation, we found out that excluding them leads to loss of information (Supplementary Figure 2, panel b). Kinex accounts for multiple phosphorylation sites within a sequence and optionally for phospho-priming, and generates all possible sub-sequences.

#### Scoring matrix

The scoring matrix represents a database of pre-calculated scores of 82,755 known phosphorylation sites (+-7 amino acids from the phospho-acceptor) curated by Johnson *et al.* Kinases are stored on columns and each row represents a phosphosite. In order to make the software efficient, we ordered each column in ascending order (the order of substrates is lost, though it is on purpose because the essential information needed here is the distribution of scores for each kinase). Given any kinase, we can quantify how likely it will phosphorylate an input substrate considering all 82k phosphosites as possible targets.

#### Phospho-acceptor favorability

Johnson *et al.*[1] used phosphorylation assays with 208 recombinant Ser/Thr kinases and substrate peptides containing either serine or threonine phospho-acceptors (as seen in their Extended Data Fig. 6 a). They also used a statistical approach to determine the phospho-acceptor favorability by summing the serine or threonine scores across all positions and scaled the results using the maximum value of the two. They found good correlation between the experimental and statistical approach (as detailed in Johnson *et al.*'s Extended Data Fig. 6 b[1]).

To score the serine/threonine phospho-acceptor favorability, we implemented the statistical method proposed by Johnson *et al.* in their Supplementary note 1.

$$\text{Sum}_S = \sum_{p=-5}^{4} M_{S,p}; \quad \text{Sum}_T = \sum_{p=-5}^{4} M_{T,p} \tag{1}$$

$$S_{\text{ctrl}} = 0.75 * \text{Sum}_S - 0.25 * \text{Sum}_T; \quad T_{\text{ctrl}} = 0.75 * \text{Sum}_T - 0.25 * \text{Sum}_S \tag{2}$$

$$S_0 = \frac{S_{\text{ctrl}}}{\max(S_{\text{ctrl}}, T_{\text{ctrl}})}; \quad T_0 = \frac{T_{\text{ctrl}}}{\max(S_{\text{ctrl}}, T_{\text{ctrl}})} \tag{3}$$

#### Kinase inference

Kinex scores each phosphopeptide and records the top 15 kinases, compiles a list of unique kinases from the identified top kinases, and determines the frequency of each unique kinase's occurrence in up-regulated, down-regulated, and un-regulated phosph-peptides. Finally, it constructs a 2 × 2 contingency table for each kinase based on its occurrence in the different regulation states:

**Table 1:** Contingency Table

| | Hit | Total |
|---|---|---|
| Upregulated | upreg_hit | upreg_total |
| Unregulated | unreg_hit | unreg_total |

Where:

- upreg_hit represents the number of sequences that each kinase phosphorylates.
- upreg_total represents the total number of upregulated sequences in the study.
- unreg_hit represents the number of unregulated sequences that each kinase may phosphorylate.
- unreg_total represents the total number of unregulated sequences in the study.

One-sided exact Fisher's test is applied along with the Benjamini–Hochberg correction. The plotting can omit the B-H correction.

The same contingency table structure is also used for down-regulated sequences, with 'downreg_hit' representing the number of down-regulated sequences that each kinase phosphorylates, and 'downreg_total' representing the total number of down-regulated sequences in the study.

Kinex stores for each sequence the top 15 upstream kinases and output them in a data frame for reporting purposes.

For kinase inference, Kinex needs the complete phosphoproteomics data-set, which includes all phosphopeptides detected without any heavy filtering. For example, if only the up- and down-regulated peptides are given, one can increase the threshold of the log transformed fold change in order to get some background (un-regulated/unchanged) information.

**Kinase activity inference** Kinex first categorises input phosphorylation sequences as down-regulated, up-regulated, or not substantially regulated (un-regulated for short), based on a user-defined threshold of fold-change. Next, Kinex identifies the highest ranked kinases for each phosphopeptide, compiles a list of unique kinases, and counts their occurrences in each regulatory category. With these results, Kinex constructs two $2x2$ contingency tables for each kinase (one table for up-regulated and un-regulated sites, and another for down-regulated and un-regulated sites). It applies the Fisher's exact test to ascertain the predominant direction of regulation.

The computational time of kinase inference for one comparison (i.e. treatment compared with control) in a phosphoproteomics dataset is less than 1 minute on a modern laptop with Intel i5 CPUs and 8GB memory.

Though sequences with multiple phosphorylation residuals increases computation time, even for the extreme cases where all sequences have 5 phosphorylation sites, we observe a comparable computational time (Supplementary Figure 1 panel c).

**A reference atlas of drug-induced kinase activity profiles** With Kinex, we pre-computed the enrichment tables of 900 samples originating from 31 drugs tested across 13 cell lines. Each sample, in this context, is uniquely represented by the combination of a drug, its concentration, the duration of treatment, the cell line, and the numbering of replicates. The database allows Kinex users to compare inferred causal kinases profiles with known drugs tested in multiple cell lines in a concentration- and time-dependent manner.

To determine the relationships between an input sample and the reference drugs, Kinex calculates Euclidean distances from the input kinase activity change profile to all 900 samples. Users can visualise the result using multiple dimensionality reduction techniques, including UMAP (Uniform Manifold Approximation and Projection), MDS (Multidimensional Scaling), and t-SNE (t-Distributed Stochastic Neighbor Embedding).

#### Validation of Kinex's ability to recover perturbed kinases

In order to validate the ability of Kinex to infer causal kinases, we curated and re-analysed four phosphoproteomics datasets with defined single perturbations[3,4,5,6]. The datasets were generated by diverse groups with multiple human cell lines. In each case, cells are treated with either control or a single treatment such as pharmacological or genetic perturbation or an external stimuli such as ionizing radiation.

In all cases, Kinex was able to recover the causal kinases and downstream mechanisms associated with the known perturbation (Supplementary Figure 2). While *Kinase Library* is also able to infer these causal kinases, we notice that the distribution of p-values returned by Kinex shift toward left compared with *Kinase Library*, suggesting that Kinex is more sensitive in detecting causal kinases (Supplementary Figure 2). Further analysis showed that it is because Kinex models phosphopeptides containing multiple phosphosites explicitly while *Kinase Library* filter such sequences. It highlights the value of modelling phospho-priming effects explicitly.

#### The Ang2/VEGF phosphoproteomics experiment

**Cell culture** Primary HUVECs (Lonza) were thawed at P6 and cultured in growth medium (Lonza, EBM™-2 + SingleQuots® kit + 2% FBS). Upon reaching confluence, cells were split onto 150 mm dishes.

**Starvation** When confluent, growth medium was changed to starving medium (Lonza, EBM™-2, 0.5% FBS).

**Stimulation** After 24 hours of starvation, cells were stimulated for 20 minutes with either 300 ng/ml Ang2, 5 ng/ml VEGF, a combination of 300 ng/ml Ang2 + 5 ng/ml VEGF, or left unstimulated for control.

**Cell lysis and storage** Post-stimulation, cells were washed once with PBS at RT (room temperature) and lysed in 500 ul of urea lysis buffer (1% SDS, 8 M urea, 50 mM ammonium bicarbonate, complete w/o EDTA, PhosSTOP). Lysates were directly put onto dry ice and stored at -80 °C until phospho-proteomics analysis.

**Sample preparation and TMT labeling** After digestion, samples were split into 2 batches: full proteome samples and phospho-proteome enriched samples. For each batch, one sample per condition was processed at the same time in a 10plex TMT run, and consistent tags were used for the same samples in each run.

**Mass spectrometry and data normalisation** Mass Spec measurements were mapped, annotated and normalised with Proteome Discoverer (PD, Thermo Fisher Scientific). Intensities for the reporter ions were summed up for each peptide and further summed to protein level for full proteome samples. For normalisation, all peptide abundances per sample were summed up and scaled by the sample with the maximum value. After export from PD, an additional reference sampled based TMT-run normalisation was performed (internal reference scaling (IRS)[7]). For each TMT-run, 2 reference samples were available (pools of all samples). Finally, data was log transformed.

**Protein and phosphopeptide quantification** In total 7540 unique proteins were quantified of which 2664 are contained in the phosphopeptide data-set. 8514 unique phospho-peptides are quantified containing 6020 phosphosites.

**Quality control and differential phosphorylation analysis** A PCA based quality control analysis was performed on IRS normalised data, including an outlier check[8]. For differential phosphorylation analysis, the log transformed phospho-peptide abundances were first adjusted for protein level abundance per sample to exclude false positive hits due to differential protein abundance. Then, linear models were fit on the adjusted data-set using limma[9] and all treatment groups were compared to the control. Resulting p-values were adjusted for FDR (False Discovery Rate)[10] per contrast. All computations were performed in R (R Core Team 2019).

**Sample aggregation and phosphosite filtering** All contrasts have been merged in one sheet, keeping the p-values and log transformed fold change. The modified sequence format is +-7 amino acids from the phospho-acceptor. When a phosphosite appears in several rows (due to its presence in several peptides), it is kept only once. Finally, the best p-value among the occurrences and its corresponding $\log_2$ fold change are assigned to the unique phosphosite.

**Phospho-proteomics analysis with Kinex** We report the result of each treatment in one separate table sheet (Supplementary Table 3). Each table contains detected modified sequences and corresponding log2-transformed fold changes. We run Kinex with phospho-acceptor favorability set to `True` and a log-transformed fold change threshold of 1. Given that there is only one site per sequence, there was no need to account for phospho-priming.

#### Rationales to test Kinex with mouse data

Several lines of evidence support the notion that the specificity of a large proportion of kinase-substrate relationships are conserved between human and other species. Sequence and structure-based analysis of specificity determinants in eukaryotic protein kinases suggests that the conservation of kinase specificity is generally high between human and mouse[11]. Computational analysis of kinase-substrate pairing across species suggests that the conservation of the pairwise specificity between human and mouse can be as high as 80%[12]. Among 77 kinase-inhibitor drugs that are approved by FDA as of September 2023, we found evidences for 49 (64%) drugs that support a likely conserved kinase-substrate relationship between human and mouse orthologues (Supplementary Figure S5). However, few existing tools for kinase inference have been explicitly tested with phosphoproteomics data from mouse studies. Therefore, we attempted to test Kinex in such settings.

#### Visualisation

Kinex employs *Plotly Express* to generate visualisations of the inference results. Kinases are colored by their phylogenetic relationships. The plot can both be interactively viewed or saved as a static image.

### Supplementary figures

#### Supplementary figure 1

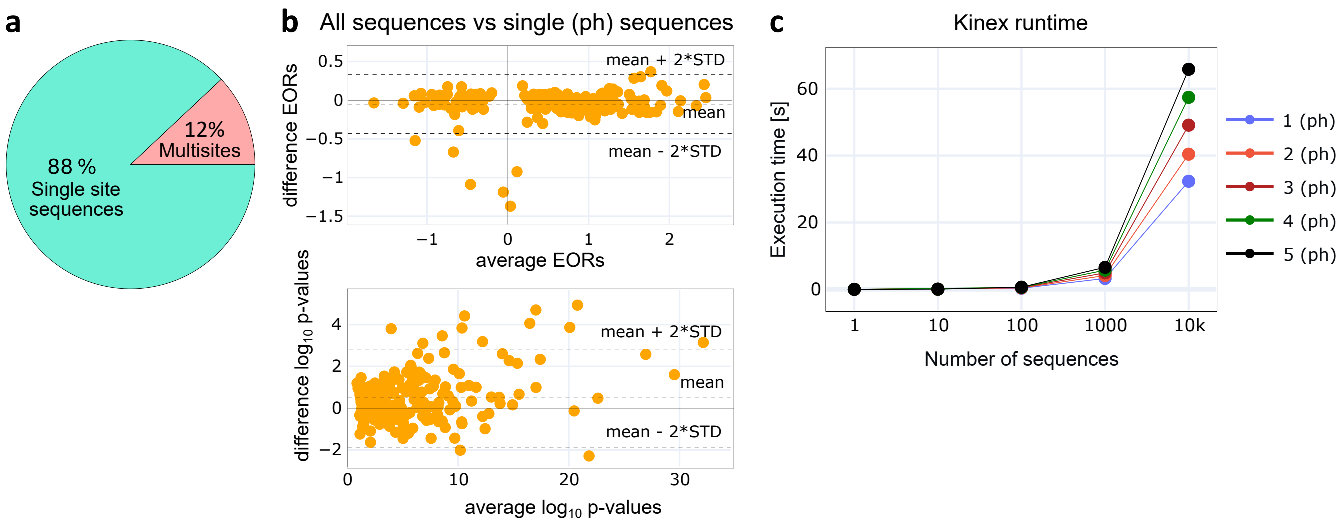


**Supplementary Figure 1: Multi-site scoring and software runtime.a**, average percentage of multi-site phosphorylation sequences across 58 experiments from Zecha *et al.***b**, M-A plot scomparing enrichment odds ratios and log10-transformed p-values inferred from two variants of analysis applied to the same dataset, which contains 13% multisite sequences. In one analysis, we used all sequences (including those with multiple phosphorylation sites), while in the other one, we filterd out multisite sequences and keep only sequences with maximum one phosphorylation site. **c**, the runtime of Kinex when applied to varying numbers of phosphorylation sequences (from 1 to 10,000) and phosphorylation sites (from 1 to 5). The values represent the averages of 10 iterations for each respective combination.

### Supplementary figure 2

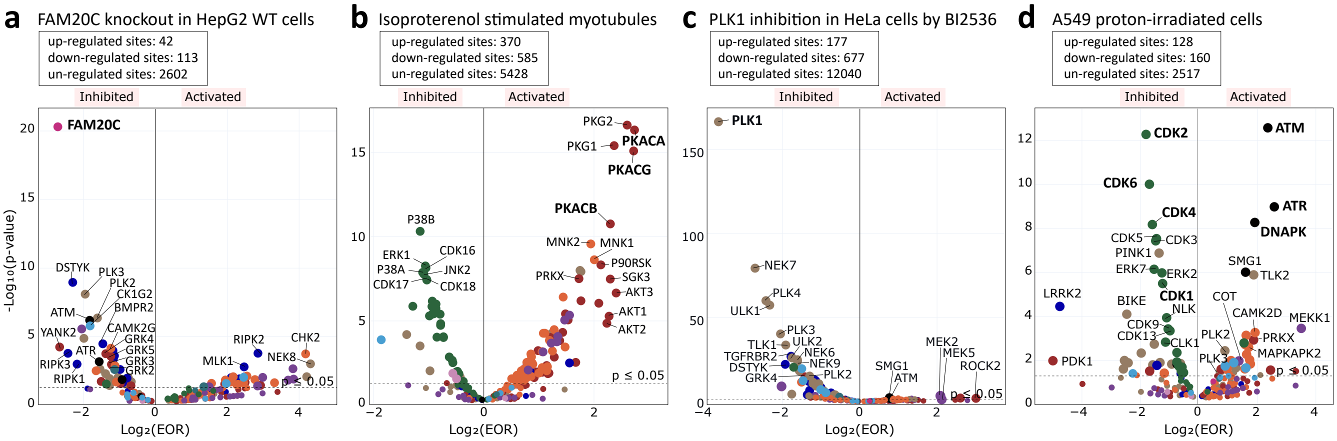


**Supplementary Figure 2: Software validation with public data-sets**. We applied Kinex to several published phosphoproteomics datasets, and observe that it is able to recover known causal perturbations. For all analysis, if not otherwise specified, we used a log2 fold-change threshold of 1.5, the scoring method "max" and we included the phospho-acceptor favorability.**a**, HepG2 cells with Fam20C knocked out with CRISPR in HepG2 cells compared with wild-type cells. We used the processed data from Phosphinator[3].**b**, L6 Myotubes after 30 min treatment with 2 $\mu$M isoproterenol[4].**c**,HeLa cells after 30 minutes treatment with 0.1 $\mu$M BI2536, a potent and selective PLK inhibitor. We used the processed data from the cells arrested at metaphase (BI mitosis)[5].**d**, A549 cells after 2h exposure to 6 Gy of proton radiation[6]. For this analysis, we used a log fold change threshold of 0.5 in order to compared with the results reported by Johnson *et al.*

### Supplementary figure 3

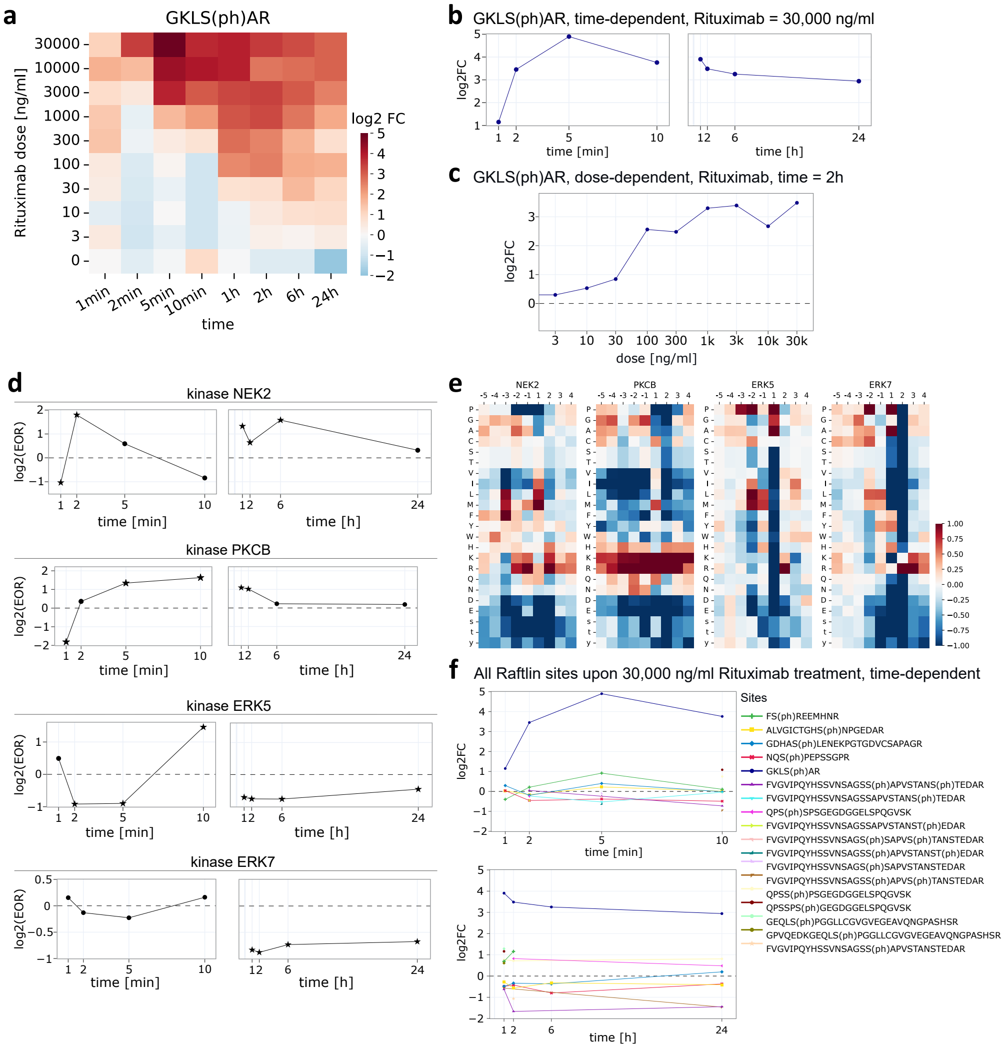


**Supplementary Figure 3: Dose- and time-dependent phosphorylation of Raftlin protein upon treatment with the monoclonal antibody Rituximab.a,**heatmap of the phosphorylation changes over dose and time at site Ser467 (GKLS(ph)AR), with color gradient and intensity indicating the magnitude of phosphorylation: blue for low magnitude, red for high magnitude and white for non-significant changes. The x-axis indicate the treatment time, from 1 minute to 24 hours, while the y-axis indicate the concentration of Rituximab that was used**b,**time-dependent phosphorylation changes at site Ser467 upon treatment with 30,000 ng/ml of Rituximab. Phosphorylation at this site increases during the first 5 minutes, reaches a maximum, and then starts to decrease.**c,**concentration-dependent phosphorylation changes at site Ser467 after 2 hours treatment with Rituximab. The magnitude of phosphorylation increases with the concentration.**d,**temporal profiles of enrichment odds ratio (EOR) profiles of the top four kinases detected by Kinex to phosphorylate the Ser467 site. For the substrate scoring we use the complete supported amino acids sequence of Ser467, namely SKGKLS(ph)ARDK, as shown in Fig.1e.**e,**the heatmaps represents the log transformed normalised and scaled PSSM scores for the kinase motifs of NEK2, PKCB, ERK7, and ERK5. Each cell in the heatmap corresponds to a specific amino acid at a specific position in the motif. The gradient and intensity of each cell indicates the PSSM score: dark red colors for higher scores (a higher preference for the amino acid occurring at that position) and dark blue color for lower scores (a lower preference for the amino acid occurring at that position)**f,**temporal changes in phosphorylation at all sites of Raftlin protein upon treatment with 30,000 ng/ml of Rituximab. Ser467 is the only site among the present Raftlin sites in the study that is sustainably and substantially up-regulated. Each line on the plot represents a phosphorylation site sequence. In all line graphs, dash lines indicate $x = 0$; asterisks represent the enrichment odds ratio values for which $p \leq 0.05$; circles are used for enrichment odds ratio values for which $p > 0.05$.

#### Supplementary figure 4

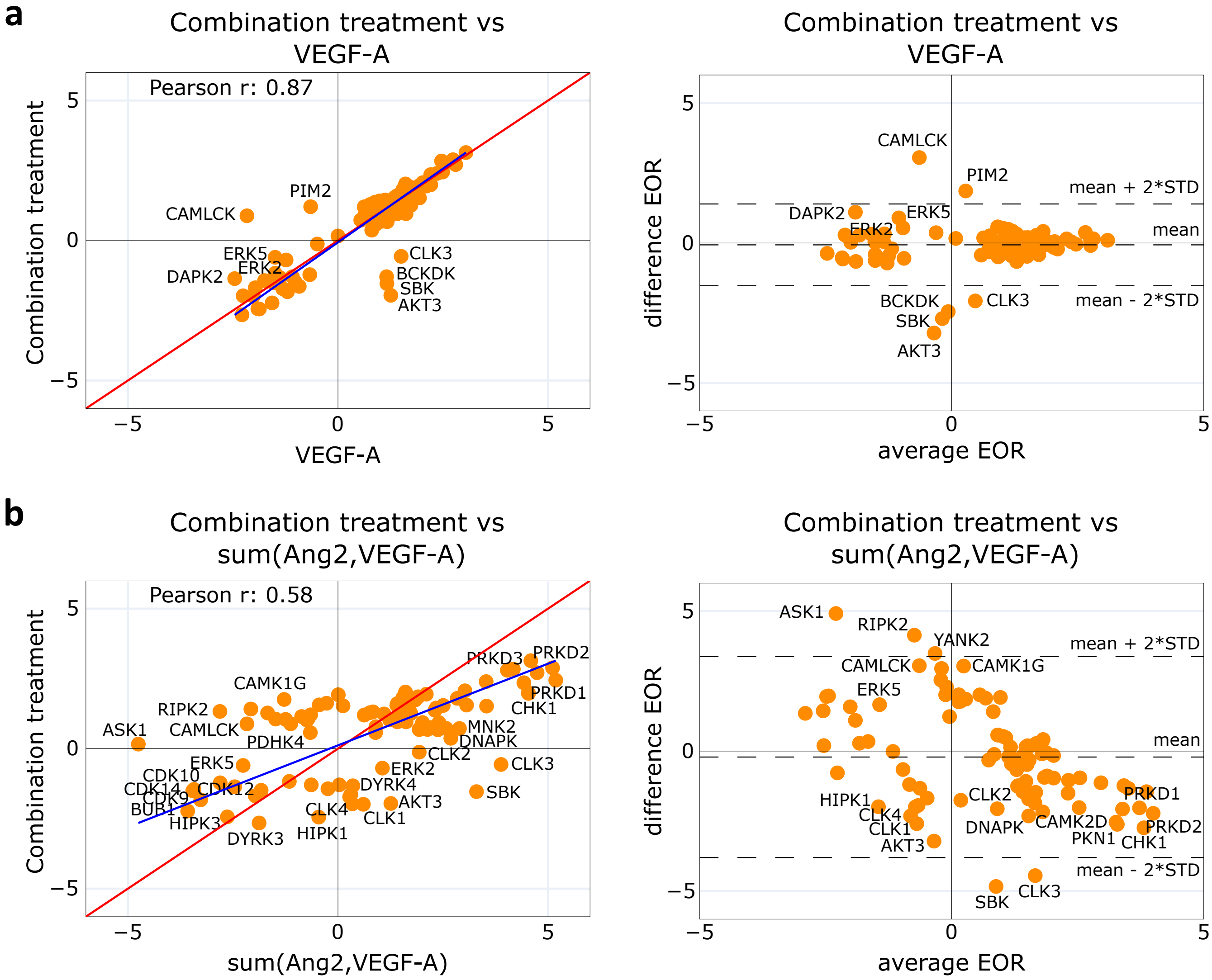


**Supplementary Figure 4: Comparison of treatments effects of growth factors on HUVECs.** Scatter plot and MA plot comparing the combination treatment of Ang2 and VEGF-A against: **a**,the treatment with VEGF-A alone or **b**,the cumulative enrichment odds ratio of cells treated separately with Ang2 and VEGF-A. For all the plots, the enrichment odds ratio value was obtained by running Kinex with the same settings as described in Fig.2. Left panels show the scatter plots, where the blue line represents the ordinary least square (OLS) regression trend line, the red line represents the line of perfect fit, and the Pearson correlation coefficient is also indicated. On the right are the MA plots, where the difference between the inferred activities of the two conditions are plotted on the y-axis, while the arithmetic mean is plotted on the x-axis. Solid black lines indicate $x = 0$ and $y = 0$. Dash lines indicate twice the standard deviation of all points, assisting spotting outliers.

#### Supplementary figure 5

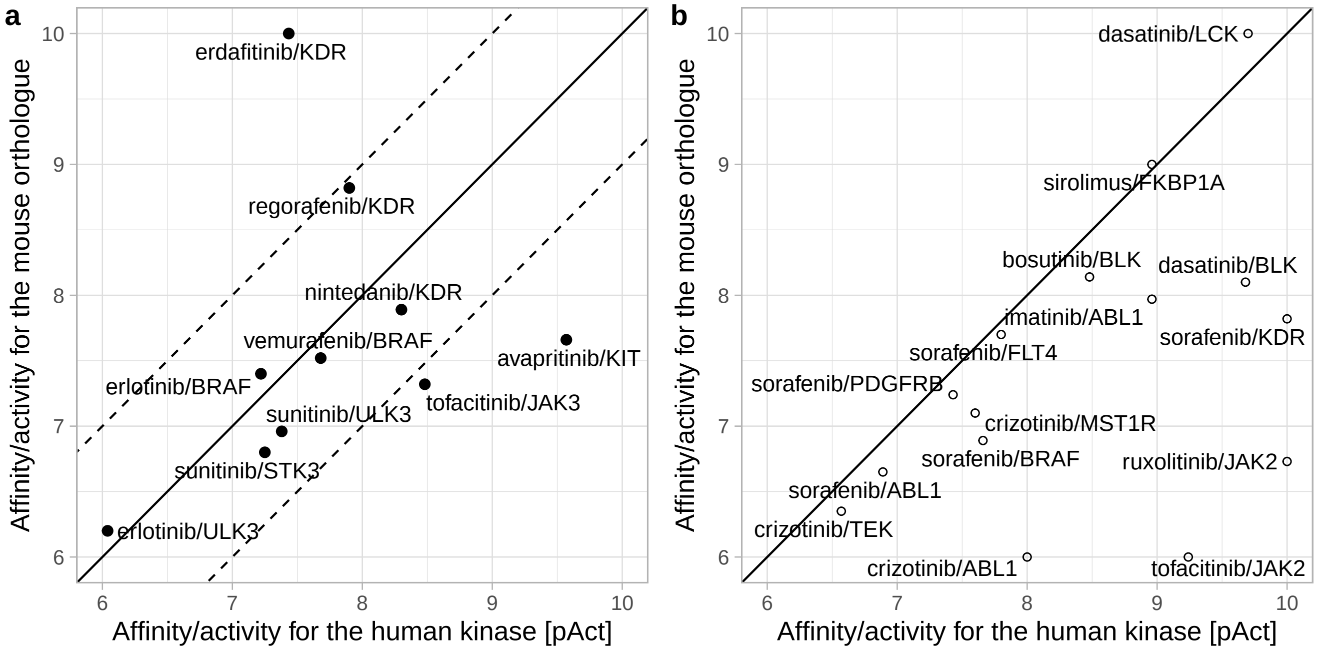


**Supplementary Figure 5: Conservation of kinase specificity between human and mouse for FDA-approved inhibitors**. **a**: Binding affinity (Kd) or inhibition (IC50) of the human kinase (x-axis) and the mouse orthologue (y-axis) by the inhibitors. Different units (either Kd or IC50) are used between kinase/inhibitor pairs, however for each pair, the same unit is reported for human and mouse kinases. Either Kd or IC50 values are transformed into the pAct value, which is defined by the absolute log10-transformation of the original value. The solid line indicates the same potency/inhibition against two orthologues, and dash lines indicate the difference of one log(10) between the two species. **b**: Similar to the left plot, though this time the type of measurement (Kd, Ki, or IC50) is different between human and mouse orthologues. Since measurements differ between the two species, it makes little sense to compare the values quantitatively. However, notice that for most cases, inhibitors that bind preferentially with human kinases (pAct>=6, or IC50, Ki, or Kd < 1mM) bind, albeit less strongly, with mouse kinases. Kinase/inhibitor binding data were curated from DrugCentral 2023.

#### Supplementary figure 6

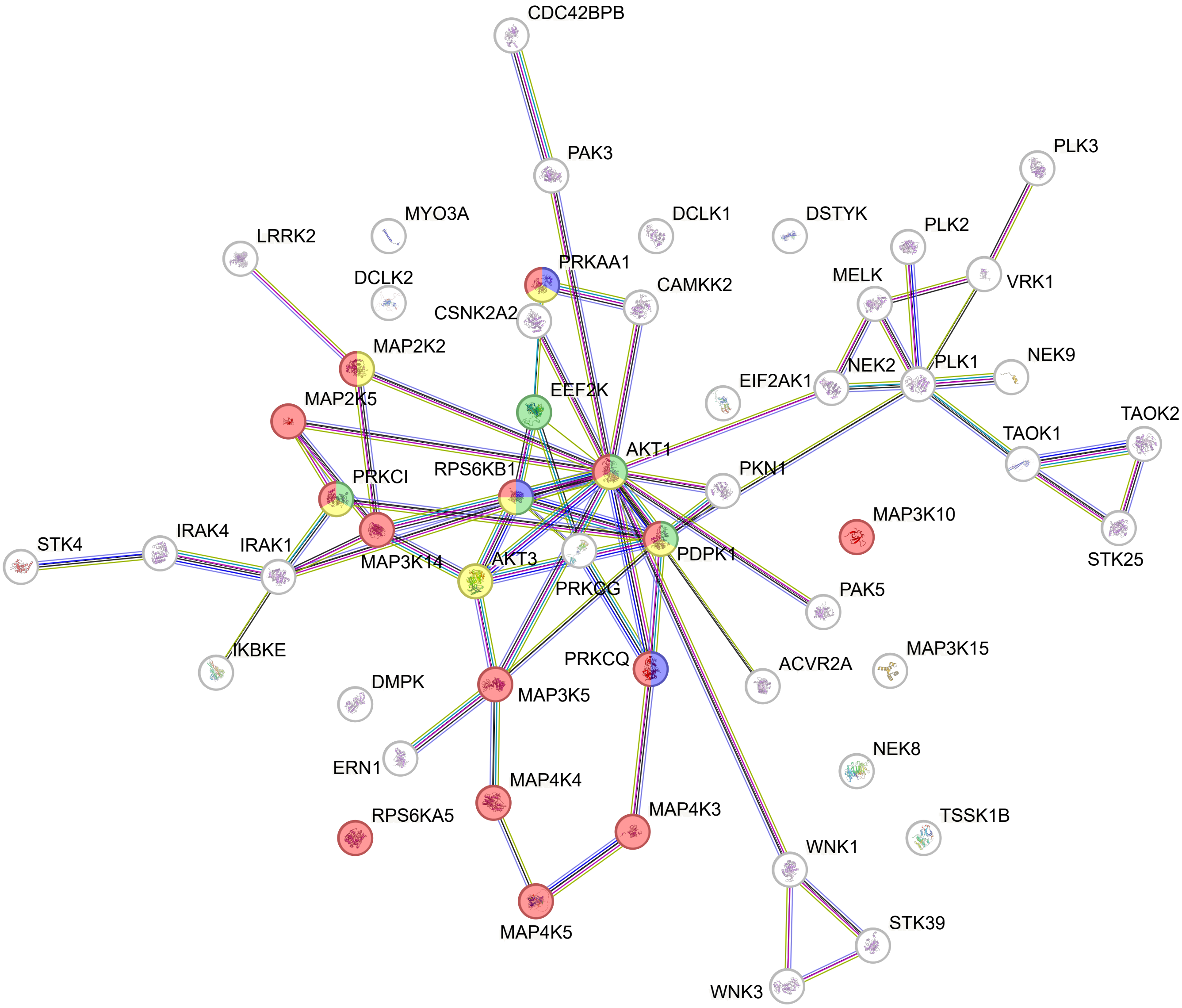


**Supplementary Figure 6: STRING pathway analysis of top inferred modulated kinases detected by Kinex in insulin signaling.** We queried STRING with the top enriched kinases, using the Uniprot identifier as input (Supplementary Table XX). Color-highlighted nodes were identified as part of the insulin signaling pathway: red, yellow and green nodes indicate known positive regulators, while blue nodes indicate a known negative regulator. The edges represent the protein-protein interactions, with the color of the edges indicating the type of evidence: blue for co-occurrence, black for co-expression, purple for experimental evidence, green for neighborhood evidence, and light blue for database evidence.

**Supplementary figure 7**

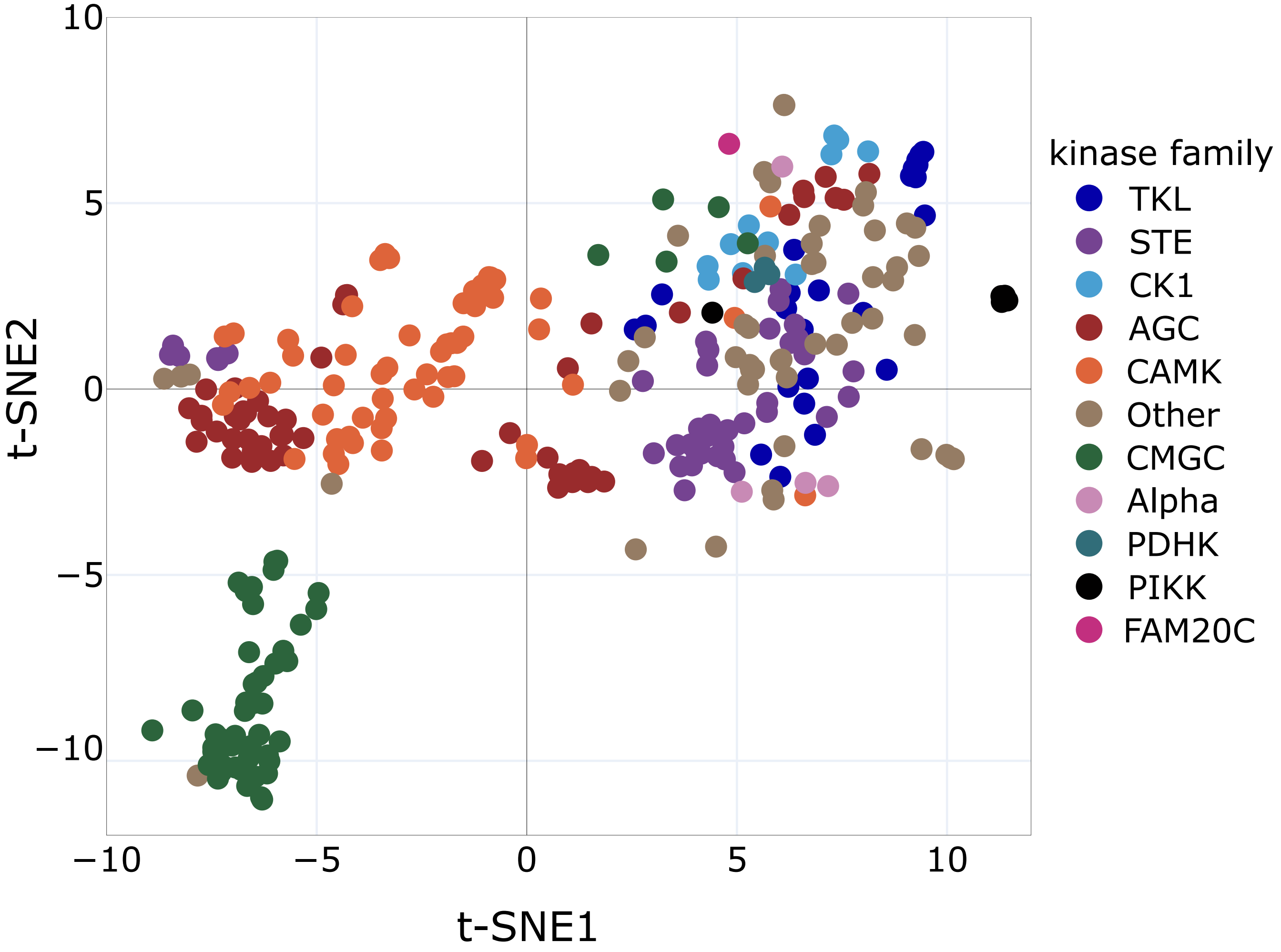


**Supplementary Figure 7: Kinase motifs landscape.** The t-SNE plot shows the diversity of the serine and threonine kinase motifs. Each point on the map represents a kinase colored by its family. The spatial proximity between the points reflects the similarity between their recognition motifs. The plot was generated using the normalised and scaled PSSM scores. We use the cosine similarity for the dissimilarity measure and ran t-SNE implemetned in the *sklearn* library with $perplexity = 50$ and $random - state = 0$.

#### Supplementary figure 8

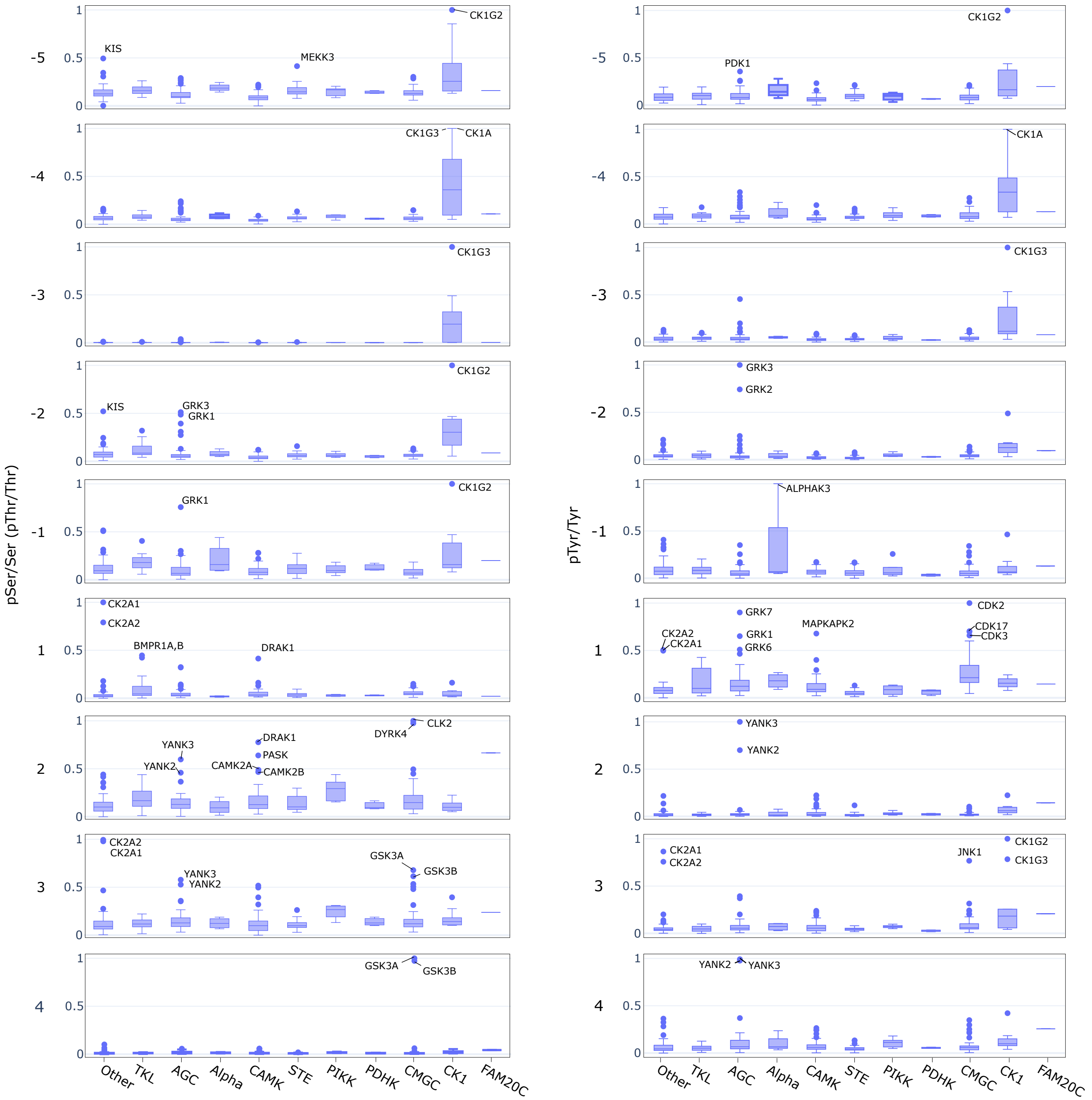


**Supplementary Figure 8: A boxplot of position-specific phospho-priming preferences across kinase families.** The ratio between between phosphoresidues and amino acids (pThr/T, pSer/S, pTyr/Y) was computed using the normalised and scaled scores from PSSM tables provided by Johnson **et al.** Points indicate outliers beyond 1.5 times of the interquantile ranges. The x-axis indicates kinase families. The y-axis represents the ratio between the phosphoresidue and amino acid which have been scaled to a [0,1] range using sklearn's `MinMaxScaler` function. Each subplot from top (-5) to bottom (4) represents the relative position of the phosphoresidue/amino acid to the phospho-acceptor.
